# Structural Interpretation of Hydrogen-Deuterium Exchange with Maximum-Entropy Simulation Reweighting

**DOI:** 10.1101/769398

**Authors:** R.T. Bradshaw, F. Marinelli, J.D. Faraldo-Gómez, L.R. Forrest

## Abstract

Hydrogen-deuterium exchange combined with mass spectrometry (HDX-MS) is a widely applied biophysical technique that probes the structure and dynamics of biomolecules in native environments without the need for site-directed modifications or bio-orthogonal labels. The mechanistic interpretation of measured HDX data, however, is often qualitative and subjective, owing to a lack of quantitative methods to rigorously translate observed deuteration levels into atomistic structural information. To help address this problem, we have developed a methodology to generate structural ensembles that faithfully reproduce HDX-MS measurements. In this approach, an ensemble of protein conformations is first generated, typically using molecular dynamics simulations. A maximum-entropy bias is then applied *post-hoc* to the resulting ensemble, such that averaged peptide-deuteration levels, as predicted by an empirical model of a value called the protection factor, agree with target values within a given level of uncertainty. We evaluate this approach, referred to as HDX ensemble reweighting (HDXer), for artificial target data reflecting the two major conformational states of a binding protein. We demonstrate that the information provided by HDX-MS experiments, and by the model of exchange, are sufficient to recover correctly-weighted structural ensembles from simulations, even when the relevant conformations are observed rarely. Degrading the information content of the target data, e.g., by reducing sequence coverage or by averaging exchange levels over longer peptide segments, reduces the quantitative structural accuracy of the reweighted ensemble but still allows for useful, molecular-level insights into the distinctive structural features reflected by the target data. Finally, we describe a quantitative metric with which candidate structural ensembles can be ranked based on their correspondence with target data, or revealed to be inadequate. Thus, not only does HDXer facilitate a rigorous mechanistic interpretation of HDX-MS measurements, but it may also inform experimental design and further the development of empirical models of the HDX reaction.

**Statement of significance:** HDX-MS experiments are a powerful approach for probing the conformational dynamics and mechanisms of proteins. However, the mechanistic implications of HDX-MS observations are frequently difficult to interpret, due to the limited spatial resolution of the technique as well as the lack of quantitative tools to translate measured data into structural information. To overcome these problems, we have developed a computational approach to construct structural ensembles that are maximally diverse while reproducing target experimental HDX-MS data within a given level of uncertainty. Using artificial test data, we demonstrate that the approach can correctly discern distinct structural ensembles reflected in the target data, and thereby facilitate statistically robust evaluations of competing mechanistic interpretations of HDX-MS experiments.

## Introduction

Upon exposure to a deuterated solvent such as D_2_O, labile hydrogen atoms present in protein side chains and backbones will readily exchange for deuterium. The rate of this process is influenced by the chemical features of the exchanging groups and by conditions such as pD or temperature, and is also critically dependent on protein conformation (1, 2). Consequently, measurements of Hydrogen-Deuterium exchange (HDX) rates are increasingly used as a direct probe of protein dynamics. Moreover, by combining HDX with mass spectrometry (HDX-MS), this approach has become feasible also for large complexes and membrane proteins, even at low concentrations (3).

Typically, HDX-MS is carried out using so-called bottom-up and continuous labeling strategies, in which proteins are deuterated for varying amounts of time, quenched, proteolytically fragmented, and purified in the solution phase, before analysis of the individual peptide fragments by mass spectrometry. For each identified fragment, typically 5-20 residues in length, deuterium incorporation is then reported as the change in peptide mass over time. Because side-chain and terminal-amine deuterons exchange back relatively rapidly with protons during analysis, HDX-MS data reports exclusively on backbone-amide exchange. This ability to directly probe protein dynamics has led to diverse applications (4), including studies of allostery (5–7), epitope mapping for protein-protein or protein-lipid interactions (8–11), effects of ligand binding (12–15), mechanisms of membrane proteins (16–22) and dynamics of large macromolecular complexes (23–26). This progress notwithstanding, the interpretation of HDX-MS data in structural and mechanistic terms has been, generally speaking, largely qualitative and lacking objective metrics.

No matter the protein system, interpretation of HDX-MS data requires an understanding of the processes reflected in the exchange kinetics. For any given backbone amide under a given set of conditions (pH, temperature, etc.), the most rapid rate of exchange occurs when the residue is in a completely unstructured, solvent-accessible conformational state of the protein. Under these circumstances, the value of the intrinsic exchange rate constant, 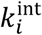 for residue *i*, is determined predominantly by steric and electronic effects from neighboring sidechains (27, 28). In a folded conformational state, by contrast, amides will be partially or fully occluded from solvent and/or engaged in hydrogen bonding. This structural protection can diminish the intrinsic rate constant by several orders of magnitude. In this case, exchange is better described as a two-step process: first, a structural transition must occur from a so-called non-competent exchange state to a competent one; this step is followed by the intrinsic chemical exchange reaction with rate constant 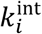 (2, 29). If the structural transition entails only local alterations rather than complete unfolding, an equilibrium between the exchange-competent and non-competent states may be reached rapidly, even more so than the hydrogen-deuterium substitution; this situation is referred to as occurring with ‘EX2’ kinetics. The overall exchange rate under these conditions is thus given by the product of the equilibrium constant for the structural transition and the intrinsic rate, 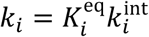. This relationship is commonly expressed as 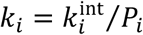, where *P*_*i*_ denotes the ‘protection factor’ for each amide, which in turn relates to the free energy difference between the non-competent and competent states, Δ*G* = *RT* ln *P*_*i*_. Following these concepts, HDX data is commonly interpreted in terms of the degree of protein structural flexibility and solvent accessibility for a given amide.

In practice, HDX-MS experiments measure deuteration averaged over lengthy peptide fragments rather than at the single-residue level. Even with robust methods to derive high-resolution protection factors directly from experimental data (30, 31), interpretation of the observed data in structural terms is not straightforward. Oftentimes HDX levels are color-coded and mapped on known protein structures, which allows an intuitive visualization of the results and highlights dynamic or solvent-exposed protein regions. However, this kind of qualitative visual analysis can easily lead to a biased interpretation of the experimental data (32). Moreover, HDX data reflect the properties of an ensemble of protein conformations and in some cases, therefore, might not be explained by a single structural state. To address these issues, previous studies have relied on molecular simulation methods. A typical approach is to first generate a conformational ensemble for the protein of interest with molecular dynamics (MD) or Monte Carlo simulations. Empirical models that predict protection factors *P*_*i*_ for individual protein structures are then used to compute peptide deuteration levels, whose ensemble averages are correlated with the experimental data (33–43). An important caveat of this seemingly straightforward approach is that, for many cases of interest, a simulation may not accurately represent the conformational ensemble reflected by the experimental data, for example due to force-field inaccuracies or incomplete sampling. Thus, even if a perfectly accurate empirical model for *P*_*i*_ was used, the predicted protection factors might deviate substantially from measured data.

Here, we develop and test a simulation methodology to construct conformational ensembles that faithfully reflect a given set of target HDX-MS data, for a given empirical model of *P*_*i*_. This approach, which we refer to as HDX ensemble reweighting (HDXer), is based on concepts outlined in previous studies and applied to other types of biophysical data (44–54), but not yet to HDX-MS. In brief, our approach uses a maximum-entropy criterion to reweight the configurations obtained computationally *post-hoc*, so that calculated ensemble-averaged peptide deuterated fractions reproduce measured values, within a given level of uncertainty. This approach aims to adjust populations in a heterogenous conformational ensemble such that they conform ideally to the experimental data, while taking into account all potential sources of uncertainty. Importantly, this method allows us to rank the correspondence between a given HDX-MS dataset and several candidate conformational states, based on the degree of bias required to reproduce the experimental results.

To evaluate the validity of HDXer, we analyze artificial HDX-MS data generated for the two major conformations of TeaA, the accessory substrate-binding protein of the bacterial TRAP membrane transport system (55). This protein undergoes a large-scale conformational change upon substrate binding, which has previously been characterized through extensive enhanced-sampling simulations (55). To rigorously assess the performance of our method we reweight these simulation data so that calculated deuteration levels match a set of artificial HDX-MS data that reflects pre-defined populations of the two conformational states. We then ask whether the conformations favored by the reweighting indeed correspond to the structural states used to generate the target data. To explore the reliability of the method in practical applications, we also performed the reweighting for target data whose information content had been progressively degraded, either by increasing the length of the protein fragments or by reducing the sequence coverage. Encouragingly, the results show that the proposed approach always recovers the key features of the correct structural ensemble, even when sparse HDX data are targeted.

## Theory and Methods

### Calculation of HDX residue protection factors and peptide deuterated fractions

To predict deuterium uptake based on structural snapshots (obtained from MD simulations or other molecular modeling method), we first calculate the protection factor for each residue *i, P*_*i*_, using the method of Best and Vendruscolo (33). Specifically, the free-energy difference between exchange-competent and non-competent states of a residue is approximated by a linear function of the numbers of H-bonds and heavy-atom contacts of the corresponding backbone amide, denoted as *N*_H,*i*_ and *N*_C,*i*_, respectively:

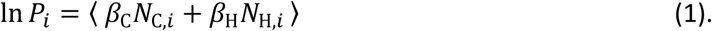

The notation ⟨… ⟩ signifies an ensemble average over all snapshots available. *N*_C,*i*_ is calculated as the number of non-hydrogen atoms within 6.5 Å of the amide N atom of residue *i*, excluding atoms in residues *i*–2 to *i*+2; *N*_H,*i*_ is the number of O or N atoms within 2.4 Å of the amide hydrogen atom. In the original formulation by Best and Vendruscolo, the scaling factors *β*_C_ and *β*_H_ are set to 0.35 and 2.0, respectively. These values reflect an empirical optimization with respect to experimental HDX data for several water-soluble proteins (33); however, their optimal value depends on the protein or experimental conditions (36), and therefore we will treat them as optimizable parameters.

In addition to *P*_*i*_, we consider the intrinsic exchange rate constant for each residue type, 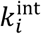, from Bai and coworkers, updated for acidic residues and glycine (27, 28). Deuterated fractions for peptide segments of the protein, 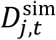, can then be calculated for any given timepoint of exchange, *t*, using the exchange rate constants of each individual residue and according to first-order kinetics. That is:

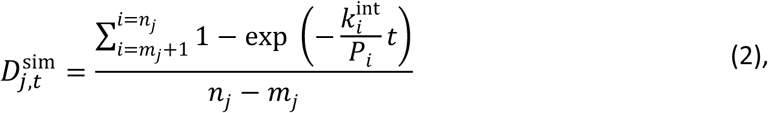

where *m*_*j*_ and *n*_*j*_ are the first and last residue numbers of the *j*-th protein fragment respectively. Note that proline residues do not have an exchangeable amide proton and were therefore excluded from the deuterated fraction calculation. The first residue (*m*_*j*_) in each peptide segment was also omitted from the average, as hydrogens in the amine N-terminus are labile after proteolytic fragmentation and are assumed to have fully exchanged back to protons during the HDX-MS purification and analysis step. It should also be noted that in direct comparisons of experimental and predicted data, the measured deuterated fractions should be corrected for the fraction of D_2_O/H_2_O in the reaction buffer and for back-exchange during the analysis process. Both corrections can be achieved by normalizing to deuterated fractions observed in identical control experiments performed under maximal deuteration conditions (32).

### Maximum-entropy ensemble reweighting with HDX data

In this section we describe the basic formulation for calculating corrections to the statistical weight of the individual structural snapshots in an ensemble, each denoted by **X**_*k*_, such that the ensemble-averaged deuteration fractions reproduce a set of HDX experimental data. Our approach is related to that of Marinelli & Fiorin (46), in which the only bias applied is that strictly required to conform to the experiments, following the so-called maximum-entropy principle (44, 45, 53, 54, 56). In general terms, the minimal bias needed to correct the mean value of one or more observables of interest is provided by a linear function of those observables, added as a perturbation term to the molecular force field or energy function, *U*(**X**) (44). In this case, the target observables are *P*_*i*_ (or functions thereof) (Eq. 1-2), and therefore the corrected forcefield is defined as:

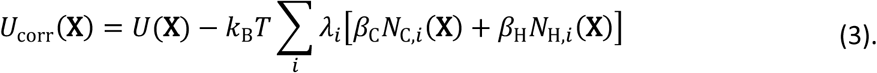

In the initial sample, the statistical weight of each configuration **X**_*k*_ is proportional to exp{− *U*(**X**_*k*_)/*k*_B_*T*}. Similarly, in the corrected ensemble these weights are proportional to exp{− *U*_Corr_(**X**_*k*_)/*k*_B_*T*}. The set of weight adjustments we seek, Ω(**X**_*k*_), are therefore simply a Boltzmann factor of the linear term of Eq. 3:

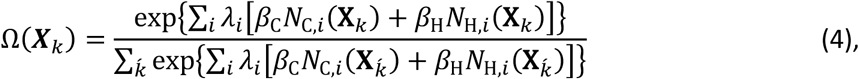

in which the denominator is a normalization term calculated by summing over all simulation configurations.

The scaling factors *λ*_*i*_ in Eq. 3-4 are the key adjustable parameters in this methodology. These parameters will be uniquely determined so that deuteration fractions deduced from the re-weighted ensemble fit the experimental data within a defined level of uncertainty, *ρ*_err_, and with the smallest possible bias. To quantify this bias, we report the amount of apparent work, *W*_app_, required to reweight the ensemble. In formal terms, the optimal value of *λ*_*i*_ is at the global minimum of the following (Kullback-Leibler) likelihood function (46, 57):

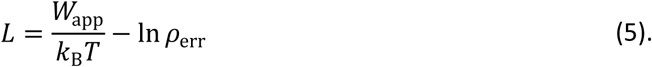

The apparent work, *W*_app_, depends on the correction to the potential applied in Eq. 3 as follows:

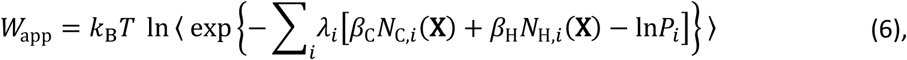

where ⟨… ⟩ denotes a mean value over the corrected ensemble, or in other words, a weighted average according to the weights of Eqn. 4. Note that *W*_app_ is related to the Kullback-Leibler divergence between the initial and corrected ensembles (46, 57, 58), *D*_KL_ = *W*_app_/*k*_B_*T* = ∑_*k*_ Ω(**X**_*k*_) ln Ω(**X**_*k*_) + ln *N*, where *N* is the number of simulation frames.

The function *ρ*_err_ is an error distribution that, for simplicity, we assume to be Gaussian:

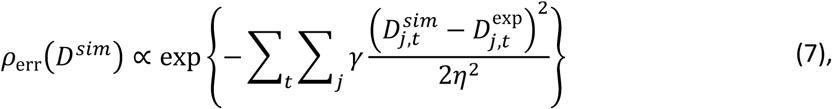

where the parameter *γ* controls the final level of agreement with the experiments (see below), *η* is an estimate of the uncertainty, and 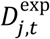 and 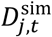 are the experimental and predicted deuterated fractions, respectively. 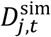 is calculated according to Eqn. 2 using the protection factors for each amide, but after adjusting for reweighting, ln *P*_*i*_ = ⟨*β*_C_*N*_C,*i*_ + *β*_H_*N*_H,*i*_⟩ = ∑_*k*_[*β*_C_*N*_C,*i*_(**X**_*k*_) + *β*_H_*N*_H,*i*_(**X**_*k*_)]Ω(**X**_*k*_).

In practice, we use a gradient-based minimization of the likelihood function *L* in Eqn. 5, in which the parameters *λ*_*i*_ are calculated iteratively according to the derivative of *L*:

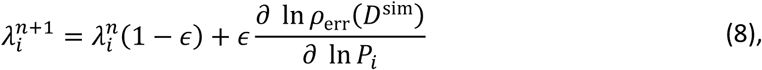

where *ϵ* is an update rate selected to ensure convergence. The weights (Eq. 4) and corrected protection factors entered into Eq. 8 depend on *λ*_*i*_ and are also updated at each iteration. The model parameters *β*_C_ and *β*_H_ are optimized at each step using a Monte Carlo procedure, to reduce the discrepancy between simulated and experimental data, measured by the mean squared deviation, MSD = *χ*^2^/*N*_D_, where 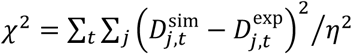 and *N*_D_ is the number of data points.

We note that although HDX-MS measures the total deuterated fraction for protein fragments, our approach uses the minimal bias condition to spread such experimental information across individual residues (Eq. 3-8). Nevertheless, if multiple experimental datapoints incorporating a single amide are available, the contribution of that amide to the ensemble correction is constrained by a simultaneous fit to all the experimental data. Therefore, in practical applications of reweighting, the inclusion of HDX-MS measurements for overlapping peptide segments will ultimately lead to enhanced resolution.

### Reweighting parameters and metrics of robustness

In the reweighting procedure, the presence of unknown errors in predicted and experimental data is considered by setting a parameter *γ* (Eq. 7) that regulates the variance in the error distribution and that can be tuned to achieve a compromise between the applied bias and the level of agreement with experiments (46, 57). To identify a reasonable value of *γ*, a decision plot of *W*_app_ *vs*. MSD can be constructed for different values of *γ*. Typically, the presence of undetermined, systematic errors, such as forward-model uncertainty or sampling inefficiency, induces a rapid increase of the work value below a certain value of MSD, resulting in an L-shaped decision plot (see **Fig. 6B**). In this case, a reasonable value of *γ* can be found at the kink of the L-curve, provided that the associated value of *W*_app_ is within a few (2-3) *k*_B_*T*.

### TeaA simulation data and generation of the artificial target HDX data

The simulation data used for ensemble reweighting were taken from the unbiased replica (∼45 ns, with frames at 1 ps intervals) of bias-exchange metadynamics simulations of TeaA performed previously (55), including both ‘closed’ and ‘open’ states of TeaA (**Fig. 1B**). The artificial HDX-MS data used as a target for the reweighting was created from this trajectory so as to represent a rapidly-interconverting conformational ensemble comprising 60% closed and 40% open states. Specifically, two reference configurations were chosen to represent typical ‘closed’ and ‘open’ states (**Fig. 1B**), and two sub-ensembles of closed and open configurations (corresponding to 37.2% and 1.6% of the initial frames, respectively) were then obtained by extracting frames in which the root-mean-square deviation (RMSD) of the Cα atoms was < 1.0 Å from those in the closed or open reference structures. The remaining 61.2% of frames remained unassigned.

**Figure 1.**
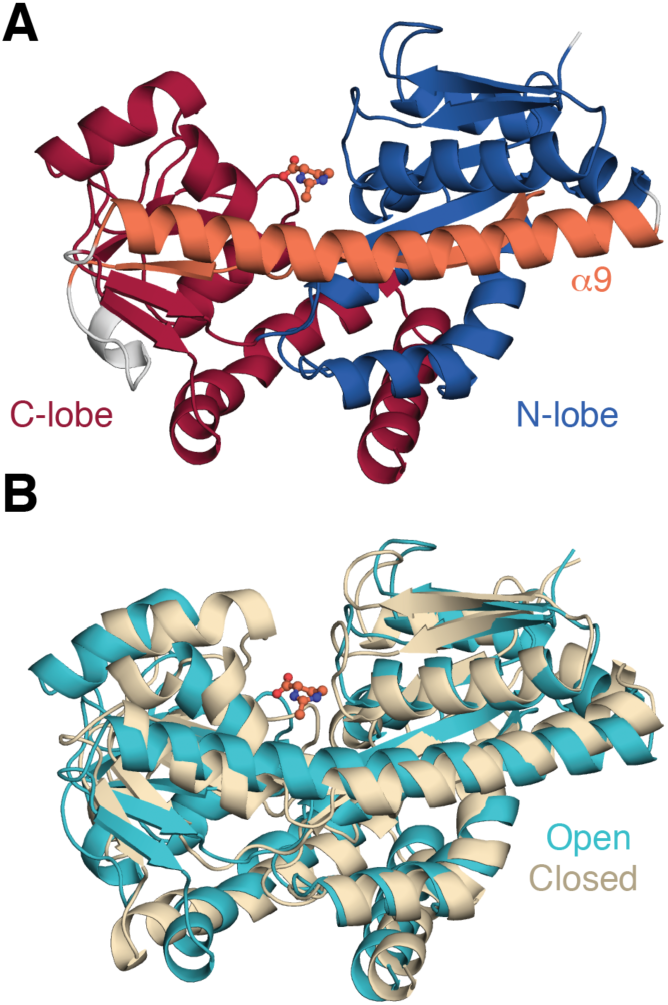
Structure of ectoine-bound TeaA in open and closed conformations. (**A**) Representative open structure shown as cartoon helices, highlighting the N-lobe (*red*), the C-lobe (*blue*), and the *β*4/α9 segments (*orange*) that span both lobes. The ectoine ligand bound to the central binding cleft is shown in ball-and-stick representation. (**B**) Overlay of representative open (*cyan*) and closed (*wheat*) conformations. The C_α_ RMSD between the two conformations is 3.2 Å.

Artificial target HDX-MS data sets were then derived from the closed and open sub-ensembles according to Eqn. 2. Residue protection factors for the mixed target ensemble were calculated as 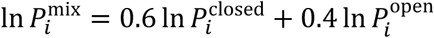, in which 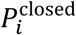 and 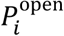 represent protection factors calculated across the sub-ensemble of the closed and open conformations, respectively. Protection factors were calculated using Eqn. 1, with *β*_C_ = 2.0 and *β*_C_ = 0.35. Artificial HDX-MS data were also constructed for the open and closed ensembles separately, using the same values of 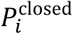 and 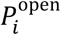. All artificial data was calculated at timepoints of 0.167, 1.0, 10.0, 60.0, and 120.0 min. These timepoints reflect typical HDX-MS experiments, and capture both short- and long-timescale EX2 exchange. A scheme of the generation and use of the artificial mixed-ensemble HDX data for ensemble refinement is provided in **Fig. S1**.

To assess the impact of segment averaging and sequence coverage upon reweighting, multiple TeaA HDX-MS datasets were generated. The largest set of artificial HDX measurements, obtained at residue-level resolution and with full sequence coverage, comprised 294 residues at 5 timepoints, for a total of 1470 individual predicted observables to be refined against. To evaluate the effect of segment averaging, five other target datasets were generated, in which TeaA was divided into fragments of size 5, 10, 15, 20 or 50-residues, including prolines. Note that, since deuteration of the N-terminal amine is excluded from HDX-MS data, neighboring protein segments were defined with a one-residue overlap (*e*.*g*., 1-10, 10-19, etc.). The final peptide segment in each dataset was extended up to the C-terminal residue 310. Analysis of the effect of sequence coverage was based on the 10-residue segment target HDX dataset, which comprises 34 peptides, from which coverage was reduced in five cumulative steps from 100% to 20% of the sequence (6-7 peptides at each step; **Fig. S2**). Assuming that buried peptides are less likely to be proteolytically hydrolyzed, we preferentially excluded peptides with lower solvent accessibility.

### Trajectory clustering

To rationalize the results of the reweighting, the structures (‘samples’) in the final ensembles were clustered based on pairwise C_α_ RMSD with the density-based algorithm DBSCAN, as implemented in scikit-learn v0.21.2 (59). The minimum size of a cluster, *n*, was set to 10% of the total ensemble size, but the contribution of each frame to *n* corresponded to the weight assigned after ensemble reweighting (Eq. 4) and normalized to the number of structures in the entire ensemble. The maximum radius, ε, which defines the neighborhood of an individual sample, was chosen by evaluating cluster quality for the ensemble obtained after reweighting to the residue-level dataset with *γ* = 10^3^. Scanning a range of values of ε from 10.546 Å to 105.46 Å (equivalent to pairwise RMSD values of 0.05 Å or 0.50 Å, respectively) on this test set revealed well-defined clusters with high silhouette scores (60) at an ε value of 42.187 Å (a pairwise RMSD of 0.20 Å).

### Data availability

All underlying data used in this study is made freely available (DOI: 10.5281/zenodo.3385169), including the initial simulation trajectories, target HDX datasets, and analysis code for extracting contacts and H-bonds, generating artificial target datasets, reweighting ensembles, and clustering. The code and underlying data used to create figures is also available in this repository.

## Results

### The TeaA test system undergoes a substantial conformational change

In the proposed computational approach, we seek to be able to reweight a heterogenous structural ensemble so that it optimally reflects a given set of HDX-MS data. The success of such a method requires that it be able to detect and up-weight the protein configurations that are most consistent with the data, but also to detect and down-weight those that are not. To meaningfully test this method, therefore, one must begin with a sample that is sufficiently heterogeneous, for a system with several states of known structure. To this end, we considered the ectoine-binding protein TeaA, and extracted a broad sample of configurations from enhanced-sampling MD simulations carried out in a previous study (55). The structure of TeaA consists of two distinct lobes interconnected by a single β-strand (β4) and a single α-helix (α9) (**Fig. 1A**). Ectoine binding at a central cleft between the lobes fosters a clamshell-like structural change (**Movie S1**), whereby the distance between lobes changes by up to ∼10 Å. We refer to the two endpoints of this conformational change as the ‘open’ and ‘closed’ states (**Fig. 1B**). These states have nearly identical secondary structure, except that closure requires local unwinding and kinking of the α9 helix, at residues K247-L249.

The existing simulations, based on Bias-Exchange Metadynamics, capture the full range of this structural change and explain how the affinity for ectoine is modulated by the conformational state of the protein (55). This data demonstrated that the closed state of TeaA becomes most favored when ectoine is bound; however, partial opening of this bound form was also observed, and found to entail a free energy penalty of only ∼2 kcal mol_-1_ (55). Accordingly, the unbiased replica in these simulations samples open, closed, and intermediate configurations of the protein (**Fig. S3**). This structural heterogeneity makes this data an ideal choice as a reference set on which to test the ability of our reweighting method.

### Artificial HDX-MS data for open and closed TeaA

To test the protocol proposed here, we need target HDX data sets for each of the conformation states of the protein. Moreover, these HDX data sets must also be distinct from each other. To our knowledge, however, no experimental HDX-MS data exists for TeaA. We therefore decided to generate artificial, high-resolution HDX data for the two major states of TeaA (open and closed), to evaluate whether a hypothetical experiment would yield a measurable contrast. To this end, we extracted separate ensembles of open and closed conformations from the simulation data and compared the predicted deuterated fractions at the single-residue level for each set (calculated using Eqn. 1-2). The HDX data was generated at single-residue resolution and across five timepoints, in order to capture both spatial and temporal differences in deuterium uptake at high resolution (see Theory & Methods).

We observed substantial differences between the deuterated fractions of closed and open states (**Fig. 2A**), confirming that these artificial data sets are well suited for our purpose. As might be expected, this contrast is most pronounced at the binding site interface and in the α9 helix (**Fig. 2B**). Interestingly, though, subtle differences are also observed across almost the entire protein and vary from one timepoint to another. These complex patterns cannot be easily interpreted visually, e.g., by mapping the data onto the representative structures (**Fig. 2B**), as they reflect the dynamical nature of the simulated ensembles. For the same reasons, such comparisons based on single structures also offer limited insights into experimentally-determined HDX-MS data, as has been noted elsewhere (32). The striking variability of the idealized artificial data for TeaA further illustrates the need for an ensemble perspective to rigorously interpret HDX measurements at the structural level.

**Figure 2.**
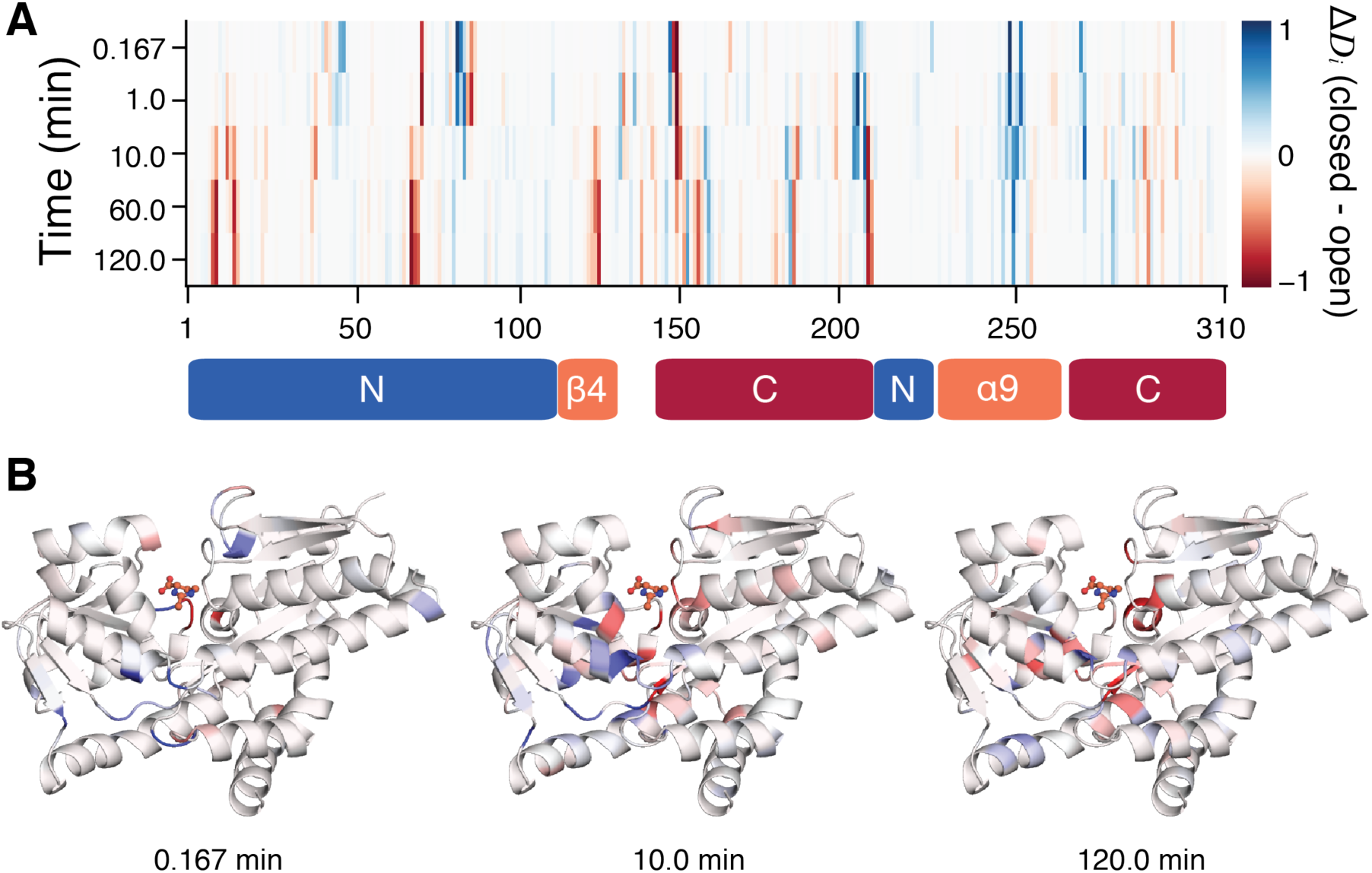
Difference in predicted deuterated fractions between closed and open ensembles of TeaA. (**A**) By-residue Δ*D*_*i*_ = *D*_*i*,closed_ − *D*_*i*,open_ for each timepoint, where red indicates that a residue is more deuterated in the open conformation than in the closed, while blue indicates the opposite. Domain definitions are indicated using bars beneath the plot. (**B**) Representative closed structure of TeaA, colored by residue Δ*D*_*i*_ at the 0.167, 10 and 120-minute timepoints. The largest Δ*D*_*i*_ values are observed for residues either lining the central binding cleft or involved in the partial unfolding of helix α9, but are clearly not uniform across timepoints.

### Ensemble reweighting with idealized single-residue HDX target data

To begin to evaluate the HDXer method, we next produced artificial HDX-MS data for a hypothetical measurement in which TeaA spontaneously interconverts between closed and open states, populating these states in a 60:40 ratio. In contrast, the sample derived from the unbiased metadynamics replica (hereafter referred to as the reference ensemble) comprises a heterogeneous set of conformations including closed, open, and unassigned (decoy) states at a ratio of 37.2: 1.6: 61.2 (see Methods). We note, however, that the decoy structures, although unassigned, do share structural similarities with either open or closed states, as demonstrated by the continuity of the RMSD distributions (**Fig. 3A, 3B**, cyan). The challenge for the HDXer method, therefore, is to identify to the appropriate weights for each and all of the configurations in the reference ensemble so that ensemble-averaged HDX levels calculated for the reweighted sample exactly reflect the 60:40 ratio of open and closed conformations in the target dataset.

**Figure 3.**
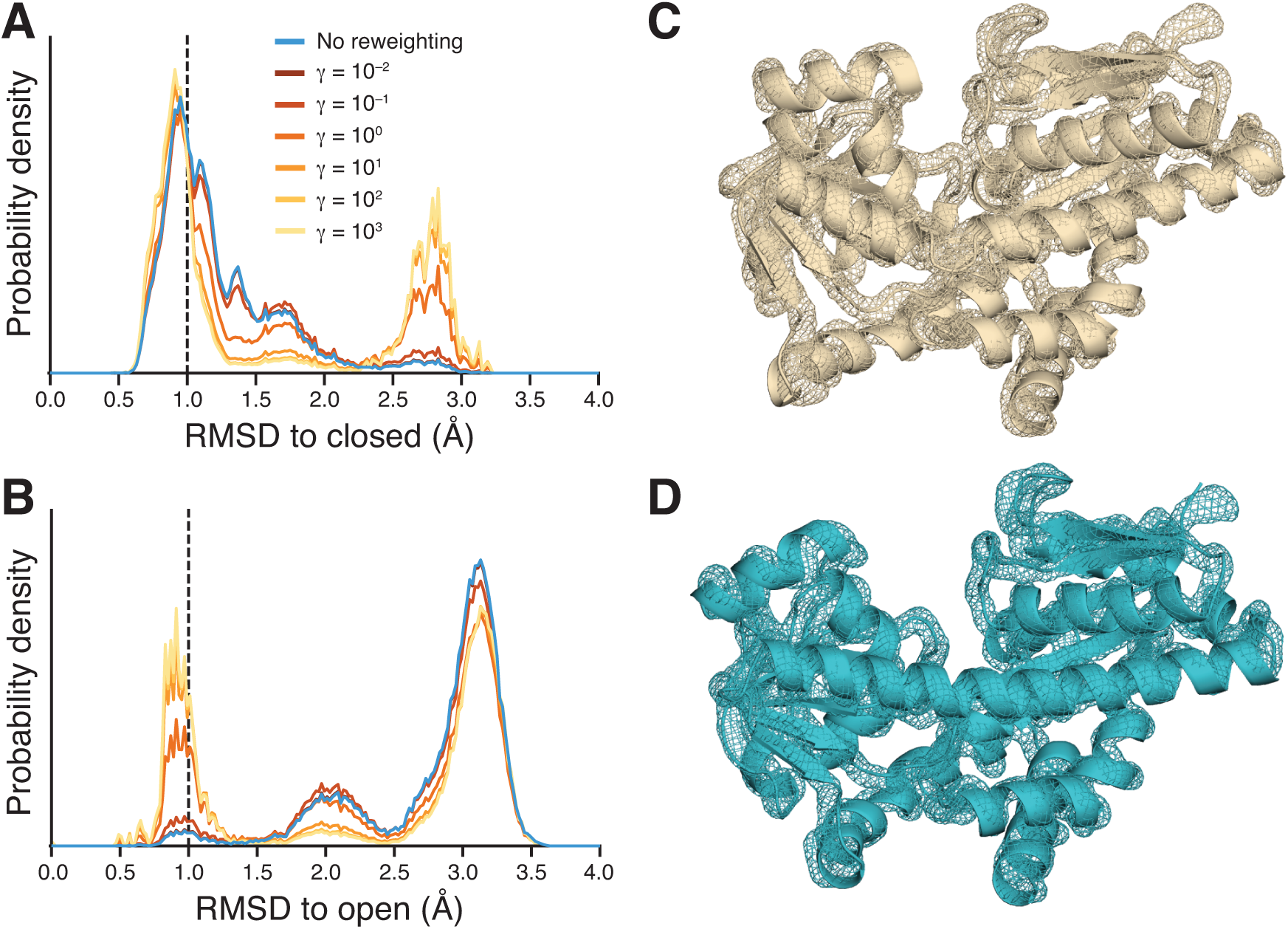
Effects of HDX ensemble reweighting at single-residue resolution. (**A & B**) Probability distributions of the RMSD with respect to the closed (A) or open (B) reference structure of TeaA for the initial, reference ensemble (cyan) and for ensembles obtained after reweighting with progressively higher *γ* values (dark brown to orange to yellow). The dashed line indicates the 1.0 Å RMSD cutoff used to define frames as belonging to the closed (A) or open (B) ensemble. (**C & D**) Ensemble density maps of the closed (C) or open (D) clusters extracted by structural clustering after reweighting with *γ* = 10^3^. The mesh reflects the density of backbone N, CA, and C atoms overlaid onto the representative closed (C) or open (D) structure of TeaA. Maps were created using the AtomProb (61) feature of Xplor-NIH v2.51, and are shown at 0.25σ.

As expected, without reweighting, the predicted HDX levels for the reference ensemble were in poor agreement with the target HDX data (MSD = 2.2 x 10^−3^), owing to the mismatch in populations. Ensemble reweighting with HDXer, however, succeeds in matching the target data (**Fig. S4, Fig. 3**). By increasing the value of the parameter *γ* in Eq. 7, an increasingly tighter agreement with the target HDX data was achieved (**Fig. S4A**), requiring a larger apparent work, *W*_app_, to be applied (**Fig. S4B**) and resulting in an increasing deviation from the initial reference ensemble. Enforcing closer agreement with the target HDX (by increasing *γ*) resulted in the gradual development of a RMSD distribution profile containing two distinct peaks, corresponding to the closed and open states of TeaA, as the decoy trajectory frames became downweighted (dark brown – orange – yellow, **Fig. 3A-B**). Notably, the bimodal features of the target distribution could already be detected with only a small applied bias of *W*_app_ = 0.9 kJ mol^−1^ relative to the 2.6 kJ mol^−1^ bias applied at *γ* = 10^3^. After reaching an MSD ≤ 10^−7^ (reweighting with *γ* ≈ 10^2^ or larger) no further substantial changes in the ensemble were observed (**Fig. 3A-B**) and *W*_app_ reached a plateau (**Fig. S4B**).

To more quantitatively characterize the outcome of the reweighting, we applied a clustering algorithm to the configurations in the reweighted ensemble obtained using *γ* = 10^3^ (see Theory & Methods). Two clusters were found: the largest clearly represented a closed conformation (**Fig. 3C**) and comprised 59.3% of the ensemble by weight, while the second cluster comprised 35.5% of the ensemble and reflected an open conformation (**Fig. 3D**). The remaining 5.2% of the ensemble consisted of outliers that, owing to structural dissimilarities and/or low weight after reweighting, could not be assigned to either of the clusters.

From the RMSD distributions of the reweighted ensembles it was clear that the final ensemble still contained a non-negligible fraction of frames > 1.0 Å RMSD to either the closed or open state. Moreover, these decoy frames were included in the extracted clusters alongside the correct frames. These observations raise concerns about the fidelity that can be achieved with ensemble-averaged observables such as these. We therefore asked how similar these decoy structures are to those in the target ensemble. According to the root-mean-square fluctuation (RMSF) of the backbone atoms, both clusters exhibited minimal structural variance, with a maximum RMSF of 1.2 Å, excluding the N-terminal residue (**Fig. S5**), and well-defined backbone density when calculated across all structures in each cluster (**Fig. 3C-D**). Therefore, the inclusion of decoy frames in fact reflected conformationally-correlated frames, indicating that the reweighting identified key structural features of the target and, based on those features, created populations of the two conformational states in good agreement with the target ratio of 60:40.

### HDX ensemble reweighting with realistic peptide segments and sequence coverage

The results so far demonstrate that HDX reweighting can successfully extract key structural features of the target ensemble using residue-resolved artificial HDX data and 100% sequence coverage. However, this level of information content is not representative of typical HDX-MS experiments, which report deuterated fractions for proteolytic fragments of a protein, while complete sequence coverage requires extensive optimization of experimental conditions. To evaluate the extent to which real-life HDX-MS data can be meaningfully interpreted with a quantitative method such as HDXer, we systematically degraded the information content of the artificial target data produced at single-residue resolution, while maintaining the 60:40 ratio of closed and open states. First, the deuterated fraction values were averaged over peptides of increasing length, from 5 to 50 residues, while maintaining full sequence coverage. Second, using fragment lengths typical for HDX-MS, sequence coverage was reduced by removing peptide segments from the target data. To compare ensembles obtained with different target datasets, for which *γ* values are not directly comparable, we instead fixed the level of agreement with the target data at MSD = 10^−6^.

Averaging the deuterated fractions over peptide segment lengths from 5 to 50 residues represents a loss of spatial resolution in the HDX-MS signal and increases the degeneracy of the structural information present in the data. When reweighting the reference ensemble, increasing the length of the segments progressively reduced the value of *W*_app_ required to achieve the same level of agreement with the target data, which is also increasingly less resolved and thus more easily reproduced (**Fig. 4A**). That is, the smaller values of *W*_app_ reflect a greater similarity between the initial and reweighted ensembles. However, this degradation of the target data translates into a reduced ability to discern between correct and decoy structures. Specifically, both the RMSD probability distributions and the structure-based clustering after reweighting (**Fig. 4B, Table 1**) show that increasing the fragment length reduced the ability of HDXer to discriminate between open and semi-open (RMSD ∼ 1.7 Å) protein structures. Indeed, using ≥ 20 residue-long segments, the semi-open state was still highly populated and was identified as a separate, unique cluster (**Table 1**). For long fragments such as these, therefore, a quantitative interpretation is not possible, unless information from overlapping (redundant) peptides is available (**Fig. S6**). Overall, however, for peptide lengths typical of HDX-MS experiments (5-20 residues), HDX reweighting correctly identified the trends in closed, open, and decoy state populations present in the target data.

**Table 1.** Cluster populations after ensemble reweighting with segment-averaged target data.

| Segment length | Closed (%) | Open (%) | Semi-open (%) | Outliers (%) |
| --- | --- | --- | --- | --- |
| 1 | 59.1 | 34.4 | – | 6.4 |
| 5 | 59.5 | 31.0 | – | 9.5 |
| 10 | 58.9 | 26.6 | – | 14.6 |
| 15 | 59.1 | 24.9 | – | 16.0 |
| 20 | 59.5 | 21.0 | 15.6 | 3.9 |
| 50 | 61.2 | 10.3 | 22.8 | 5.7 |

Populations are measured as percentage by weight of the total ensemble. Predicted deuterated fractions from the reweighted ensembles fit the target data with $MSD = 10^{-6}$ . The data for segment length = 1 represents a reweighting with residue-resolved target data.

**Figure 4.**
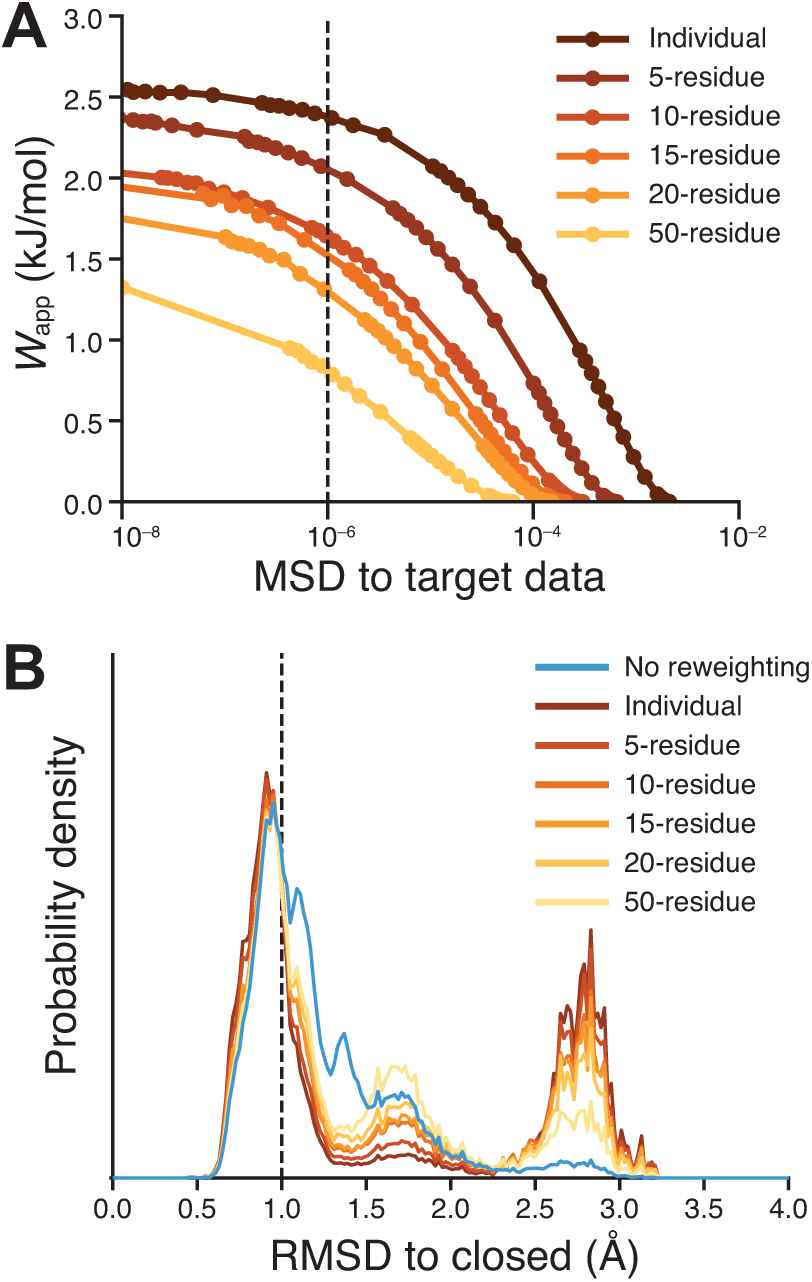
Effects of segment averaging on ensemble reweighting. (**A**) Decision plot showing the work applied during reweighting, against the MSD of the reweighted ensemble to target HDX data. Circles indicate independent reweighting experiments. (**B**) RMSD probability distributions, with reference to the closed TeaA structure, before (*cyan*) and after ensemble refinement to MSD = 10^−6^. In both panels, reference data from reweighting performed with individual residue deuterated fractions is shown in dark brown, while the data obtained by increasing the peptide segment lengths is shown using gradual color variation from light brown to orange to yellow.

Even at low levels of amide resolution, the target data up to this point covered the entire length of the protein. Loss of sequence coverage also increases the degeneracy of the structural information present in HDX-MS data. We therefore investigated the effects of reducing coverage, using the 10-residue long segment dataset analyzed earlier, for which reweighting at 100% sequence coverage resulted in cluster populations of 58.9% and 26.6% for the closed and open states, respectively (**Fig. 4B** and **Table 1**).

As expected, gradual degradation of the sequence coverage also reduced the value of *W*_app_ required to match the target data, e.g. with MSD = 10^−6^ (**Fig. 5A**), for the same reasons discussed above for the increased peptide lengths. The effect in terms of structural interpretation was also similar: reducing coverage incorrectly increased the contribution of semi-open states relative to 100% coverage (**Fig. 5B**). This effect was particularly marked when the coverage was ≤ 40% (**Table 2**).

**Table 2.** Cluster populations after ensemble refinement with target data covering smaller proportions of the protein.

| Coverage (%) | Closed (%) | Open (%) | Semi-open (%) | Outliers (%) |
| --- | --- | --- | --- | --- |
| 100 | 58.9 | 26.6 | – | 14.6 |
| 80 | 58.8 | 23.9 | 13.5 | 3.8 |
| 60 | 56.6 | 22.2 | 17.5 | 3.7 |
| 40 | 55.7 | 17.4 | 22.4 | 4.5 |
| 20 | 55.5 | 16.2 | 23.6 | 4.8 |

**Figure 5.**
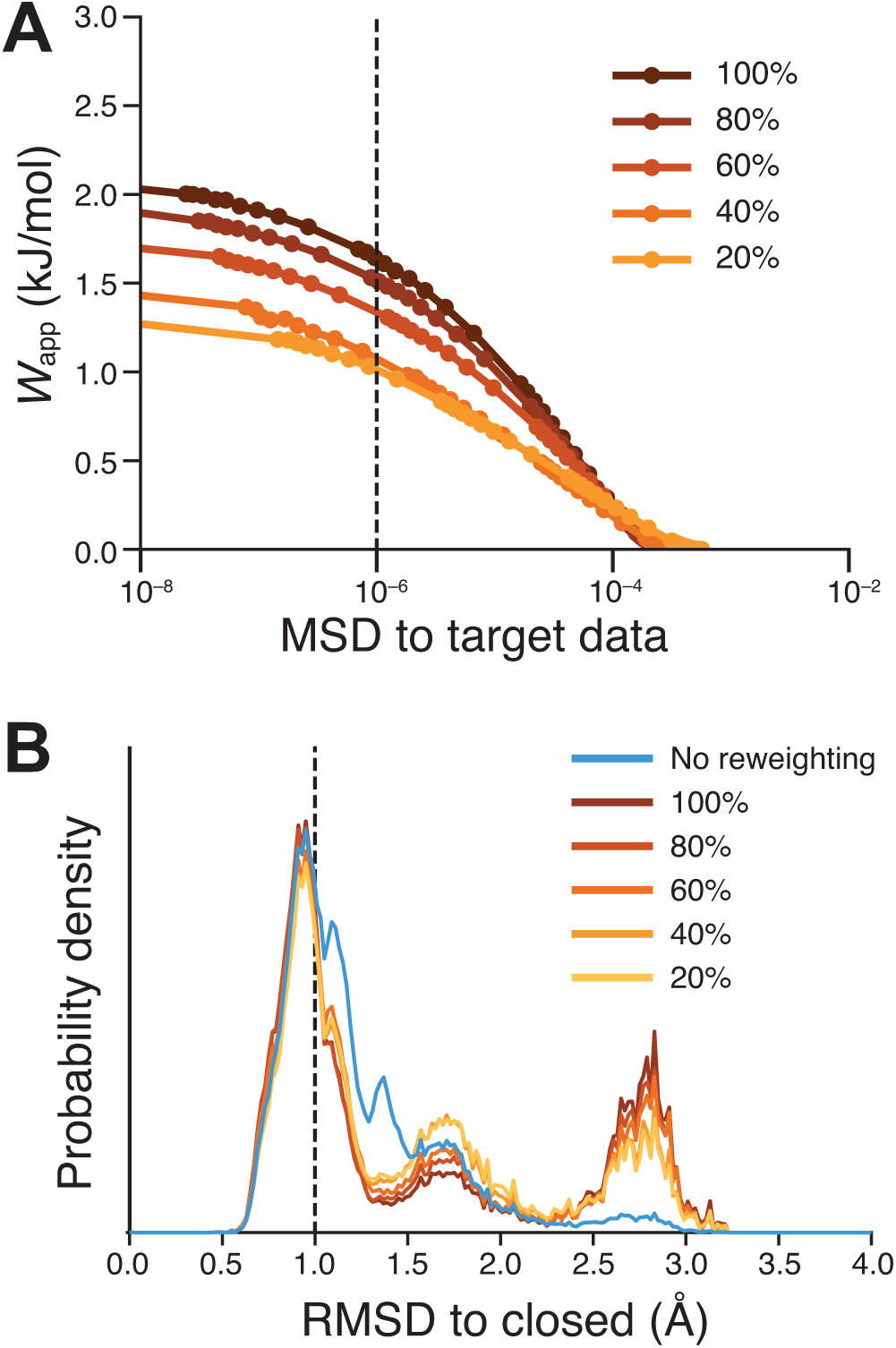
Effects of reduced sequence coverage on ensemble reweighting. (**A**) Decision plot and (**B**) RMSD probability distributions after reweighting with reduced sequence coverage in the target HDX-MS data, using target data with 10-residue segments. See legend to Fig. 4 for more details. In both panels, reference data from reweighting performed with full coverage is shown in dark brown, while the data obtained by decreasing sequence coverage lengths is shown using gradual color variation from light brown to orange to yellow.

It is perhaps surprising that even at 20% coverage, HDXer produced a 10-fold enrichment of the population of the open state, i.e., in qualitative agreement with the target data. Inspection of the peptides included in this set (**Fig. S2**) shows that at least one peptide spanning the α9 helix was included at all coverage levels. Because the conformational change in helix α9 correlates strongly with the open-to-closed transition, peptides in this helix likely include crucial target observables that allow our method to correctly discern between states of TeaA. In actual HDX experiments, this correlation might not exist for any one peptide fragment among those available, in which case 20% coverage would not likely be sufficient to derive a clear interpretation.

Overall, however, we conclude that the ability of our method to identify open and closed states from the initial sample, based only on similarity with target HDX data, does not critically depend on peptide segment length, nor does it require complete coverage. Thus, provided that the relevant conformational states are present in the reference ensemble (as has been assumed so far), HDXer will reveal the major conformational states reflected by the target HDX data, and their approximate populations, even when the HDX data is of limited spatial resolution or sequence coverage. This finding underscores the potential of the proposed method to generate structure-based interpretations of experimentally-determined HDX-MS data that are not only quantitative and objective, but also mechanistically informative.

## Discussion

Broad applicability and label-free sample preparation have made HDX-MS an increasingly attractive biophysical technique to study global biomolecular structure and dynamics under native conditions, as demonstrated by the variety of reported applications on both globular and membrane proteins and frequently-updated reviews (3, 4, 62). The major challenge, however, has been how to objectively translate the HDX data into structural information, so as to be able to derive quantitative mechanistic insights. The methodology introduced here, named HDXer, facilitates this structural interpretation by reweighting the distribution of conformations in a pre-existing ensemble, obtained for example with MD simulations, so that calculated ensemble-averaged deuteration levels match a given set of target data. Further analysis of the resulting reweighted ensemble, for example through clustering, thus provides the desired structural interpretation of the input HDX data.

As noted, the target data used to evaluate the HDXer method were artificially generated. Two factors motivated this deliberate choice. First, we aimed to focus our evaluation on the reweighting method itself, leaving aside other factors that contribute to the prediction of HDX data. By using the same empirical model of *P*_*i*_ (Eqn. 1) and EX2-like kinetics, both in the generation of the artificial HDX data and in the calculation of weights (Eqn. 4), we ensure that potential inaccuracies in this model do not influence our assessment. Similarly, by using a pre-existing configurational ensemble with a pre-defined population of states to generate the artificial data, we ensure that there is a correct answer against which our methodology can be evaluated.

The second advantage of artificial data is that it can be arbitrarily degraded, in ways that reflect the limitations of actual measurements, so as to judge the usability of the technique for structure determination. Indeed, HDX-MS studies vary greatly in terms of the level of peptide coverage and redundancy, and a priori there is no guarantee that an observed set of peptides will contain sufficient information to allow a clear structural interpretation. Our method performs optimally the better the coverage and resolution of the data, as one should expect. However, it is worth noting, and is very promising, that even with incomplete sequence coverage or lengthy peptide segments, well beyond those typically attained in well-optimized HDX-MS experiments, our re-weighting method can qualitatively identify the major conformational states contributing to the target set (**Tables 1-2**). It is also promising that estimates of the typical technical precision of HDX-MS measurements (63) suggest that achieving an ‘agreement’ with the experimental data of up to an MSD ∼ 10^−6^ (or 0.01 Da error per 10-residue peptide) is not an unreasonable goal. Our reweighting method can achieve this level of tight agreement, provided the relevant structural states are present in the initial reference ensemble, even as minority populations. Taken together, these observations lead us to conclude that HDXer will successfully provide structural insights when used to interpret actual experimental data exhibiting typical coverage and noise.

Notwithstanding these reasons for optimism, it should be noted that the ability of this or any other computational method to facilitate the interpretation of measured HDX data will depend of the fidelity of the empirical model used to calculate the residue protection factors, *Pi*, for a given structural snapshot. Our results imply that the model formulated by Best & Vendruscolo is sensitive to large conformational changes and assigns similar deuterium exchange levels to structurally correlated frames (**Fig. 3**), which are positive features well suited to ensemble reweighting. Nevertheless, no HDX prediction model has yet been shown to be uniformly accurate across different biomolecular systems (64–66), and model error could certainly lead to erroneous interpretations even after reweighting against experimental target data.

In addition to potentially uninformative data and model error, it is entirely possible that the initial reference ensemble may not include any of the structural states reflected by the HDX-MS measurements. In the case of MD sampling, this situation might arise due to sampling-time inadequacies and force-field errors. For other molecular-modeling approaches, the inherent simplifications of the energy function can be problematic. Given these different sources of potential error, it is crucial to be able to assess the reweighting process in absolute terms, i.e., to discern when the optimal solution is less than realistic. The HDXer method is equipped to do so, specifically through the calculation of the *W*_app_ required to achieve a given MSD. *W*_app_, and other metrics of reweighting robustness, such as the Kish effective sample size (51, 67), may be used to identify situations in which the relevant structural states are not part of the reference ensemble. For the TeaA model system, this situation can be exemplified by removing all the closed-state conformations from the reference ensemble before applying the HDXer method exactly as above, i.e., targeting a data set reflecting a 60% population of that state (**Fig. 6A**). The resultant decision plot (**Fig. 6B**) clearly shows that the fit cannot be improved beyond MSD ≈ 10^−4^, in contrast to the MSD < 10^−7^ attained when closed-state structures are present in the reference ensemble. Concomitantly, the *W*_app_ values increase rapidly to > 10 kJ mol^−1^ as *γ* is increased, indicating poor overlap between the reference and reweighted ensembles, i.e., only a handful of configurations have predicted HDX values in agreement with the target data. Encountering this characteristic output when using experimental data would motivate the use of enhanced-sampling methods to improve the reference pool of structures (68–71). Note that along a similar vein *W*_app_ and MSD may also be used as metrics to rank results obtained using alternative empirical models for the protection factor, so as to evaluate and improve their accuracy.

**Figure 6.**
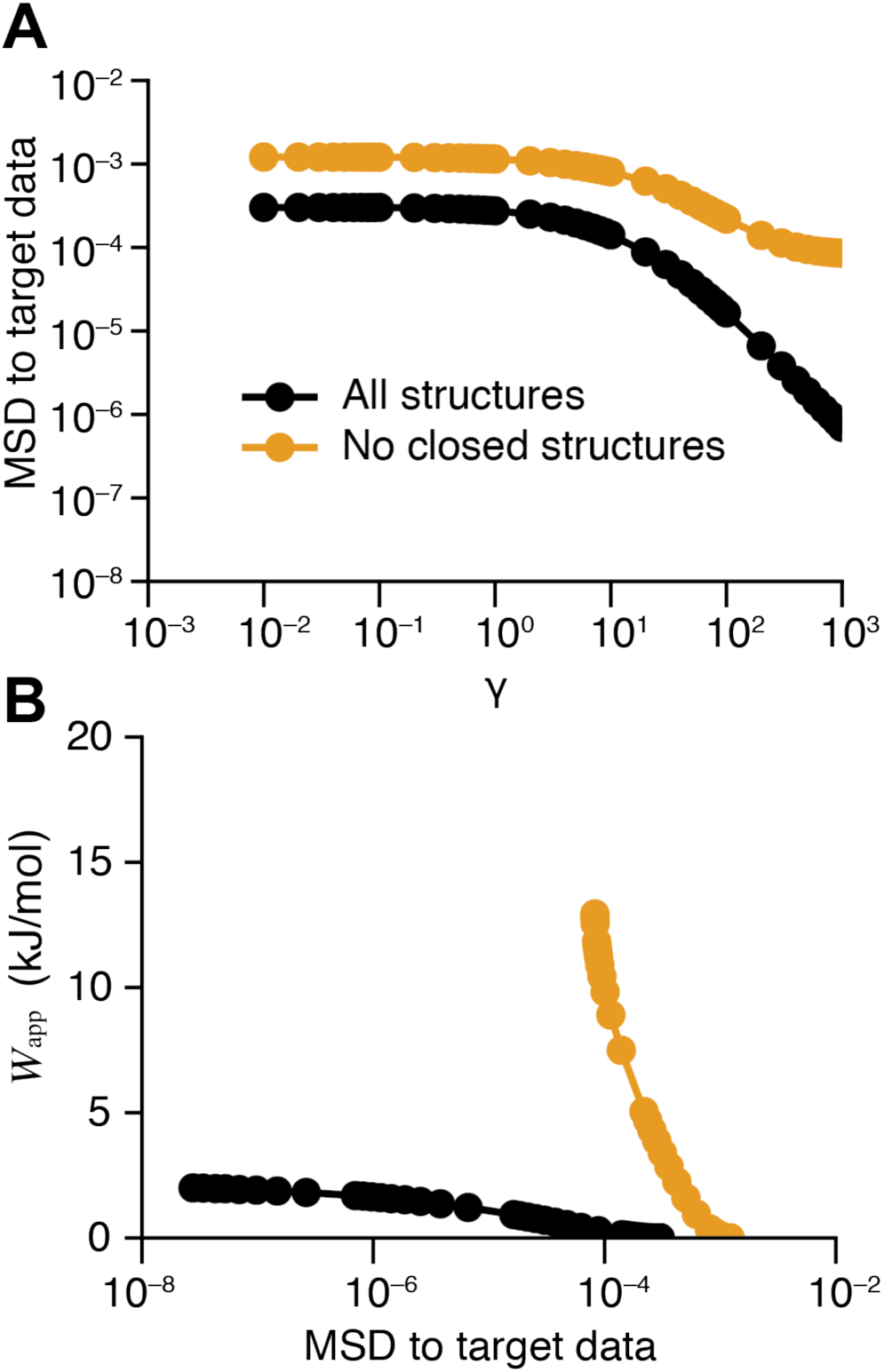
Effect of missing data on the relationship between *γ, W*_app_, and the agreement with the target data during HDX reweighting. A reference ensemble without closed TeaA structures was created by removing all frames < 1.5 Å C_α_ RMSD to the closed reference frame. Reweighting experiments were then compared between the full reference ensemble (*black*) or the degraded reference ensemble (*orange*) (**A**) The mean square deviation (MSD) between the predicted and target HDX values is consistently larger without closed TeaA structures and does not improve below MSD ≈ 10^-4^. (**B**) The lack of closed TeaA structures in the degraded reference ensemble can be identified by a rapid and substantial increase in the decision plot of *W*_app_ against MSD. Comparatively low *W*_app_ values are required to improve agreement with the target data when the reference ensemble shows adequate sampling (*black*). All reweighting was performed using the target HDX dataset with 10-residue segments.

Finally, the HDXer method could be straightforwardly applied to cross-validate the HDX data itself. Deuteration levels measured at different timepoints could be separated into training and validation sets, and inconsistencies in the resultant reweighted ensembles may reveal sources of experimental error. However, as has been extensively discussed for other ensemble refinement methods (50, 51, 57, 58, 67), disentangling the exact sources of error in a given set of reweighting results is a challenging proposition and likely to require comparison and cross-validation with multiple reference ensembles and experimental datasets.

On a technical note, it is worth underscoring that, in contrast to the canonical maximum-entropy reweighting approach, which enforces exact agreement with an experimental observable, we use a parameter *γ* to control the degree of fitness to the target – and thereby account for all uncertainties involved. Consequently, HDXer shares some of the theoretical underpinnings of Bayesian approaches used to optimally recreate experimental observables, either through ensemble reweighting or on-the-fly biased-sampling (50, 57, 72). We would argue, however, that biased sampling might not be an appropriate strategy to interpret HDX data, given the empirical nature of HDX prediction models and their imperfect correlation with experiment (64–66), and more generally, our incomplete understanding of the structural determinants of exchange across different biomolecular systems. Thus, *post-hoc* reweighting seems the most effective approach.

In conclusion, we have developed an effective maximum-entropy-based method to derive a structural-level interpretation of HDX-MS experiments *via* reweighting of conformational ensembles. We anticipate that HDXer will contribute to more systematic, quantitative analyses of HDX prediction methodologies, and aid studies of individual proteins and their functional mechanisms *via* objective structural interpretation of experimental HDX-MS measurements.

## Author contributions

Conceptualization (all authors); methodology development, programming, analysis, and writing of original drafts (RB, FM); supervision, resource and funding acquisition (JFG, LF); visualization of data (RB, FM, LF); writing - review and editing (all authors); project administration (LF).

## Acknowledgements

This research was supported by the Divisions of Intramural Research of the National Institute of Neurological Disorders and Stroke and of the National Heart, Lung and Blood Institute, National Institutes of Health, USA. This work utilized the computational resources of the NIH HPC Biowulf cluster (http://hpc.nih.gov). We are very grateful to Patrick Wintrode and Daniel Deredge for useful discussions throughout.

## References

1. Hvidt, A., and K. Linderstrøm-Lang. 1954. Exchange of hydrogen atoms in insulin with deuterium atoms in aqueous solutions. Biochim. Biophys. Acta. 14: 574–575.

2. Englander, S.W., and N.R. Kallenbach. 1983. Hydrogen exchange and structural dynamics of proteins and nucleic acids. Q. Rev. Biophys. 16: 521–655.

3. Oganesyan, I., C. Lento, and D.J. Wilson. 2018. Contemporary hydrogen deuterium exchange mass spectrometry. Methods. 144: 27–42.

4. Trabjerg, E., Z.E. Nazari, and K.D. Rand. 2018. Conformational analysis of complex protein states by hydrogen/deuterium exchange mass spectrometry (HDX-MS): Challenges and emerging solutions. Trends Anal. Chem. 106: 125–138.

5. Englander, J.J., C. Del Mar, W. Li, S.W. Englander, J.S. Kim, D.D. Stranz, Y. Hamuro, and V.L. Woods. 2003. Protein structure change studied by hydrogen-deuterium exchange, functional labeling, and mass spectrometry. Proc. Natl. Acad. Sci. U. S. A. 100: 7057–7062.

6. Klontz, E.H., A.D. Tomich, S. Günther, J.A. Lemkul, D. Deredge, Z. Silverstein, J.F. Shaw, C. McElheny, Y. Doi, P.L. Wintrode, A.D. MacKerell, N. Sluis-Cremer, and E.J. Sundberg. 2017. Structure and Dynamics of FosA-Mediated Fosfomycin Resistance in *Klebsiella pneumoniae* and *Escherichia coli*. Antimicrob. Agents Chemother. 61: e01572–17.

7. Ramirez-Sarmiento, C.A., and E.A. Komives. 2018. Hydrogen-deuterium exchange mass spectrometry reveals folding and allostery in protein-protein interactions. Methods. 144: 43–52.

8. Zhang, Q., L.N. Willison, P. Tripathi, S.K. Sathe, K.H. Roux, M.R. Emmett, G.T. Blakney, H.-M. Zhang, and A.G. Marshall. 2011. Epitope Mapping of a 95 kDa Antigen in Complex with Antibody by Solution-Phase Amide Backbone Hydrogen/Deuterium Exchange Monitored by Fourier Transform Ion Cyclotron Resonance Mass Spectrometry. Anal. Chem. 83: 7129–7136.

9. Li, J., H. Wei, S.R. Krystek, D. Bond, T.M. Brender, D. Cohen, J. Feiner, N. Hamacher, J. Harshman, R.Y.-C. Huang, S.H. Julien, Z. Lin, K. Moore, L. Mueller, C. Noriega, P. Sejwal, P. Sheppard, B. Stevens, G. Chen, A.A. Tymiak, M.L. Gross, and L.A. Schneeweis. 2017. Mapping the Energetic Epitope of an Antibody/Interleukin-23 Interaction with Hydrogen/Deuterium Exchange, Fast Photochemical Oxidation of Proteins Mass Spectrometry, and Alanine Shave Mutagenesis. Anal. Chem. 89: 2250–2258.

10. Vadas, O., and J.E. Burke. 2015. Probing the dynamic regulation of peripheral membrane proteins using hydrogen deuterium exchange-MS (HDX-MS). Biochem. Soc. Trans. 43: 773–86.

11. Martens, C., M. Shekhar, A.J. Borysik, A.M. Lau, E. Reading, E. Tajkhorshid, P.J. Booth, and A. Politis. 2018. Direct protein-lipid interactions shape the conformational landscape of secondary transporters. Nat. Commun. 9: 4151.

12. Zhu, M.M., D.L. Rempel, Z. Du, and M.L. Gross. 2003. Quantification of Protein-Ligand Interactions by Mass Spectrometry, Titration, and H/D Exchange: PLIMSTEX. J. Am. Chem. Soc. 125: 5252–5253.

13. Deredge, D.J., W. Huang, C. Hui, H. Matsumura, Z. Yue, P. Moënne-Loccoz, J. Shen, P.L. Wintrode, and A. Wilks. 2017. Ligand-induced allostery in the interaction of the *Pseudomonas aeruginosa* heme binding protein with heme oxygenase. Proc. Natl. Acad. Sci. U. S. A. 114: 3421–3426.

14. Sowole, M.A., and L. Konermann. 2014. Effects of Protein–Ligand Interactions on Hydrogen/Deuterium Exchange Kinetics: Canonical and Noncanonical Scenarios. Anal. Chem. 86: 6715–6722.

15. Masson, G.R., S.L. Maslen, and R.L. Williams. 2017. Analysis of phosphoinositide 3-kinase inhibitors by bottom-up electron-transfer dissociation hydrogen/deuterium exchange mass spectrometry. Biochem. J. 474: 1867–1877.

16. Adhikary, S., D.J. Deredge, A. Nagarajan, L.R. Forrest, P.L. Wintrode, and S.K. Singh. 2017. Conformational dynamics of a neurotransmitter:sodium symporter in a lipid bilayer. Proc. Natl. Acad. Sci. U. S. A. 114: E1786–E1795.

17. Merkle, P.S., K. Gotfryd, M.A. Cuendet, K.Z. Leth-Espensen, U. Gether, C.J. Loland, and K.D. Rand. 2018. Substrate-modulated unwinding of transmembrane helices in the NSS transporter Leu T. Sci. Adv. 4: eaar6179.

18. Möller, I.R., M. Slivacka, A.K. Nielsen, S.G.F. Rasmussen, U. Gether, C.J. Loland, and K.D. Rand. 2019. Conformational dynamics of the human serotonin transporter during substrate and drug binding. Nat. Commun. 10: 1687.

19. Nielsen, A.K., I.R. Möller, Y. Wang, S.G.F. Rasmussen, K. Lindorff-Larsen, K.D. Rand, and C.J. Loland. 2019. Substrate-induced conformational dynamics of the dopamine transporter. Nat. Commun. 10: 2714.

20. Eisinger, M.L., A.R. Dörrbaum, H. Michel, E. Padan, and J.D. Langer. 2017. Ligand-induced conformational dynamics of the Escherichia coli Na+/H+ antiporter NhaA revealed by hydrogen/deuterium exchange mass spectrometry. Proc. Natl. Acad. Sci. U. S. A. 114: 11691–11696.

21. Giladi, M., L. Almagor, L. van Dijk, R. Hiller, P. Man, E. Forest, and D. Khananshvili. 2016. Asymmetric Preorganization of Inverted Pair Residues in the Sodium-Calcium Exchanger. Sci. Rep. 6: 20753.

22. Giladi, M., L. Van Dijk, B. Refaeli, L. Almagor, R. Hiller, P. Man, E. Forest, and D. Khananshvili. 2017. Dynamic distinctions in the Na+/Ca2+ exchanger adopting the inward- and outward-facing conformational states. J. Biol. Chem. 292: 12311–12323.

23. Rostislavleva, K., N. Soler, Y. Ohashi, L. Zhang, E. Pardon, J.E. Burke, G.R. Masson, C. Johnson, J. Steyaert, N.T. Ktistakis, and R.L. Williams. 2015. Structure and flexibility of the endosomal Vps34 complex reveals the basis of its function on membranes. Science. 350: aac7365.

24. Lim, X.-X., A. Chandramohan, X.-Y.E. Lim, J.E. Crowe, S.-M. Lok, and G.S. Anand. 2017. Epitope and Paratope Mapping Reveals Temperature-Dependent Alterations in the Dengue-Antibody Interface. Structure. 25: 1391–1402.

25. van de Waterbeemd, M., A. Llauró, J. Snijder, A. Valbuena, A. Rodríguez-Huete, M.A. Fuertes, P.J. de Pablo, M.G. Mateu, and A.J.R. Heck. 2017. Structural Analysis of a Temperature-Induced Transition in a Viral Capsid Probed by HDX-MS. Biophys. J. 112: 1157–1165.

26. Radou, G., F.N. Dreyer, R. Tuma, and E. Paci. 2014. Functional dynamics of hexameric helicase probed by hydrogen exchange and simulation. Biophys. J. 107: 983–90.

27. Bai, Y., J.S. Milne, L. Mayne, and S.W. Englander. 1993. Primary structure effects on peptide group hydrogen exchange. Proteins Struct. Funct. Genet. 17: 75–86.

28. Nguyen, D., L. Mayne, M.C. Phillips, and S. Walter Englander. 2018. Reference Parameters for Protein Hydrogen Exchange Rates. J. Am. Soc. Mass Spectrom. 29: 1936–1939.

29. Hvidt, A., and S.O. Nielsen. 1966. Hydrogen Exchange in Proteins. Adv. Protein Chem. 21: 287–386.

30. Saltzberg, D.J., H.B. Broughton, R. Pellarin, M.J. Chalmers, A. Espada, J.A. Dodge, B.D. Pascal, P.R. Griffin, C. Humblet, and A. Sali. 2017. A Residue-Resolved Bayesian Approach to Quantitative Interpretation of Hydrogen–Deuterium Exchange from Mass Spectrometry: Application to Characterizing Protein–Ligand Interactions. J. Phys. Chem. B. 121: 3493–3501.

31. Skinner, S.P., G. Radou, R. Tuma, J.J. Houwing-Duistermaat, and E. Paci. 2019. Estimating Constraints for Protection Factors from HDX-MS Data. Biophys. J. 116: 1194–1203.

32. Masson, G.R., J.E. Burke, N.G. Ahn, G.S. Anand, C. Borchers, S. Brier, G.M. Bou-Assaf, J.R. Engen, S.W. Englander, J. Faber, R. Garlish, P.R. Griffin, M.L. Gross, M. Guttman, Y. Hamuro, A.J.R. Heck, D. Houde, R.E. Iacob, T.J.D. Jørgensen, I.A. Kaltashov, J.P. Klinman, L. Konermann, P. Man, L. Mayne, B.D. Pascal, D. Reichmann, M. Skehel, J. Snijder, T.S. Strutzenberg, E.S. Underbakke, C. Wagner, T.E. Wales, B.T. Walters, D.D. Weis, D.J. Wilson, P.L. Wintrode, Z. Zhang, J. Zheng, D.C. Schriemer, and K.D. Rand. 2019. Recommendations for performing, interpreting and reporting hydrogen deuterium exchange mass spectrometry (HDX-MS) experiments. Nat. Methods. 16: 595–602.

33. Best, R.B., and M. Vendruscolo. 2006. Structural interpretation of hydrogen exchange protection factors in proteins: characterization of the native state fluctuations of CI2. Structure. 14: 97–106.

34. Persson, F., and B. Halle. 2015. How amide hydrogens exchange in native proteins. Proc. Natl. Acad. Sci. U. S. A. 112: 10383–10388.

35. Wan, H., Y. Ge, A. Razavi, and V.A. Voelz. 2019. Reconciling simulated ensembles of apomyoglobin with experimental HDX data using Bayesian inference and multiensemble Markov State Models. bioRxiv, doi 10.1101/563320 (preprint posted Febr. 28, 2019)..

36. Park, I.-H., J.D. Venable, C. Steckler, S.E. Cellitti, S.A. Lesley, G. Spraggon, and A. Brock. 2015. Estimation of Hydrogen-Exchange Protection Factors from MD Simulation Based on Amide Hydrogen Bonding Analysis. J. Chem. Inf. Model. 55: 1914–1925.

37. Craig, P.O., J. Lätzer, P. Weinkam, R.M.B. Hoffman, D.U. Ferreiro, E.A. Komives, and P.G. Wolynes. 2011. Prediction of Native-State Hydrogen Exchange from Perfectly Funneled Energy Landscapes. J. Am. Chem. Soc. 133: 17463–17472.

38. Kieseritzky, G., G. Morra, and E.-W. Knapp. 2006. Stability and fluctuations of amide hydrogen bonds in a bacterial cytochrome c: a molecular dynamics study. J. Biol. Inorg. Chem. 11: 26–40.

39. Ma, B., and R. Nussinov. 2011. Polymorphic Triple β-Sheet Structures Contribute to Amide Hydrogen/Deuterium (H/D) Exchange Protection in the Alzheimer Amyloid β42 Peptide. J. Biol. Chem. 286: 34244–34253.

40. Borysik, A.J. 2017. Simulated Isotope Exchange Patterns Enable Protein Structure Determination. Angew. Chemie Int. Ed. 56: 9396–9399.

41. Liu, T., D. Pantazatos, S. Li, Y. Hamuro, V.J. Hilser, and V.L. Woods. 2012. Quantitative Assessment of Protein Structural Models by Comparison of H/D Exchange MS Data with Exchange Behavior Accurately Predicted by DXCOREX. J. Am. Soc. Mass Spectrom. 23: 43–56.

42. Petruk, A.A., L.A. Defelipe, R.G. Rodríguez Limardo, H. Bucci, M.A. Marti, and A.G. Turjanski. 2013. Molecular Dynamics Simulations Provide Atomistic Insight into Hydrogen Exchange Mass Spectrometry Experiments. J. Chem. Theory Comput. 9: 658–669.

43. Claesen, J., and A. Politis. 2019. POPPeT: a New Method to Predict the Protection Factor of Backbone Amide Hydrogens. J. Am. Soc. Mass Spectrom. 30: 67–76.

44. Pitera, J.W., and J.D. Chodera. 2012. On the Use of Experimental Observations to Bias Simulated Ensembles. J. Chem. Theory Comput. 8: 3445–3451.

45. Boomsma, W., J. Ferkinghoff-Borg, and K. Lindorff-Larsen. 2014. Combining Experiments and Simulations Using the Maximum Entropy Principle. PLoS Comput. Biol. 10: e1003406.

46. Marinelli, F., and G. Fiorin. 2019. Structural Characterization of Biomolecules through Atomistic Simulations Guided by DEER Measurements. Structure. 27: 359-370.e12.

47. Różycki, B., Y.C. Kim, and G. Hummer. 2011. SAXS Ensemble Refinement of ESCRT-III CHMP3 Conformational Transitions. Structure. 19: 109–116.

48. Cesari, A., A. Gil-Ley, and G. Bussi. 2016. Combining Simulations and Solution Experiments as a Paradigm for RNA Force Field Refinement. J. Chem. Theory Comput. 12: 6192–6200.

49. Hermann, M.R., and J.S. Hub. 2019. SAXS-restrained ensemble simulations of intrinsically disordered proteins with commitment to the principle of maximum entropy. J. Chem. Theory Comput. 15: 5103–5115.

50. Bottaro, S., T. Bengtsen, and K. Lindorff-Larsen. 2018. Integrating Molecular Simulation and Experimental Data: A Bayesian/Maximum Entropy Reweighting Approach. bioRxiv, doi 10.1101/457952 (preprint posted Oct. 31, 2018)..

51. Cesari, A., S. Reißer, and G. Bussi. 2018. Using the Maximum Entropy Principle to Combine Simulations and Solution Experiments. Computation. 6: 15.

52. Bonomi, M., G.T. Heller, C. Camilloni, and M. Vendruscolo. 2017. Principles of protein structural ensemble determination. Curr. Opin. Struct. Biol. 42: 106–116.

53. Marinelli, F., and J.D. Faraldo-Gómez. 2015. Ensemble-Biased Metadynamics: A Molecular Simulation Method to Sample Experimental Distributions. Biophys. J. 108: 2779–2782.

54. Hustedt, E.J., F. Marinelli, R.A. Stein, J.D. Faraldo-Gómez, and H.S. Mchaourab. 2018. Confidence Analysis of DEER Data and Its Structural Interpretation with Ensemble-Biased Metadynamics. Biophys. J. 115: 1200–1216.

55. Marinelli, F., S.I. Kuhlmann, E. Grell, H.-J. Kunte, C. Ziegler, and J.D. Faraldo-Gómez. 2011. Evidence for an allosteric mechanism of substrate release from membrane-transporter accessory binding proteins. Proc. Natl. Acad. Sci. U. S. A. 108: E1285–E1292.

56. Olsson, S., H. Wu, F. Paul, C. Clementi, and F. Noé. 2017. Combining experimental and simulation data of molecular processes via augmented Markov models. Proc. Natl. Acad. Sci. U. S. A. 114: 8265–8270.

57. Hummer, G., and J. Köfinger. 2015. Bayesian ensemble refinement by replica simulations and reweighting. J. Chem. Phys. 143: 243150.

58. Köfinger, J., L.S. Stelzl, K. Reuter, C. Allande, K. Reichel, and G. Hummer. 2019. Efficient Ensemble Refinement by Reweighting. J. Chem. Theory Comput. 15: 3390–3401.

59. Pedregosa, F., G. Varoquaux, A. Gramfort, V. Michel, B. Thirion, O. Grisel, M. Blondel, P. Prettenhofer, R. Weiss, V. Dubourg, J. Vanderplas, A. Passos, D. Cournapeau, M. Brucher, M. Perrot, and É. Duchesnay. 2011. Scikit-learn: Machine Learning in Python. J. Mach. Learn. Res. 12: 2825–2830.

60. Rousseeuw, P.J. 1987. Silhouettes: A graphical aid to the interpretation and validation of cluster analysis. J. Comput. Appl. Math. 20: 53–65.

61. Schwieters, C.D., and G.M. Clore. 2002. Reweighted atomic densities to represent ensembles of NMR structures. J. Biomol. NMR. 23: 221–225.

62. Masson, G.R., M.L. Jenkins, and J.E. Burke. 2017. An overview of hydrogen deuterium exchange mass spectrometry (HDX-MS) in drug discovery. Expert Opin. Drug Discov. 12: 981–994.

63. Hudgens, J.W., E.S. Gallagher, I. Karageorgos, K.W. Anderson, J.J. Filliben, R.Y.-C. Huang, G. Chen, G.M. Bou-Assaf, A. Espada, M.J. Chalmers, E. Harguindey, H.-M. Zhang, B.T. Walters, J. Zhang, J. Venable, C. Steckler, I. Park, A. Brock, X. Lu, R. Pandey, A. Chandramohan, G.S. Anand, S.N. Nirudodhi, J.B. Sperry, J.C. Rouse, J.A. Carroll, K.D. Rand, U. Leurs, D.D. Weis, M.A. Al-Naqshabandi, T.S. Hageman, D. Deredge, P.L. Wintrode, M. Papanastasiou, J.D. Lambris, S. Li, and S. Urata. 2019. Interlaboratory Comparison of Hydrogen–Deuterium Exchange Mass Spectrometry Measurements of the Fab Fragment of NISTmAb. Anal. Chem. 91: 7336–7345.

64. Skinner, J.J., W.K. Lim, S. Bédard, B.E. Black, and S.W. Englander. 2012. Protein hydrogen exchange: Testing current models. Protein Sci. 21: 987–995.

65. McAllister, R.G., and L. Konermann. 2015. Challenges in the Interpretation of Protein H/D Exchange Data: A Molecular Dynamics Simulation Perspective. Biochemistry. 54: 2683–2692.

66. Mohammadiarani, H., V.S. Shaw, R.R. Neubig, and H. Vashisth. 2018. Interpreting Hydrogen–Deuterium Exchange Events in Proteins Using Atomistic Simulations: Case Studies on Regulators of G-Protein Signaling Proteins. J. Phys. Chem. B. 122: 9314–9323.

67. Rangan, R., M. Bonomi, G.T. Heller, A. Cesari, G. Bussi, and M. Vendruscolo. 2018. Determination of Structural Ensembles of Proteins: Restraining vs Reweighting. J. Chem. Theory Comput. 14: 6632–6641.

68. Marinelli, F. 2013. Following easy slope paths on a free energy landscape: the case study of the Trp-cage folding mechanism. Biophys. J. 105: 1236–1247.

69. Peacock, R.B., J.R. Davis, P.R.L. Markwick, and E.A. Komives. 2018. Dynamic Consequences of Mutation of Tryptophan 215 in Thrombin. Biochemistry. 57: 2694–2703.

70. Camilloni, C., and F. Pietrucci. 2018. Advanced simulation techniques for the thermodynamic and kinetic characterization of biological systems. Adv. Phys. X. 3: 1477531.

71. Markwick, P.R.L., R.B. Peacock, and E.A. Komives. 2019. Accurate Prediction of Amide Exchange in the Fast Limit Reveals Thrombin Allostery. Biophys. J. 116: 49–56.

72. Bonomi, M., C. Camilloni, A. Cavalli, and M. Vendruscolo. 2016. Metainference: A Bayesian inference method for heterogeneous systems. Sci. Adv. 2: e1501177.

