## Supplementary Figures 1-6 for "Structural Interpretation of Hydrogen-Deuterium Exchange with Maximum-Entropy Simulation Reweighting"

### Contents

**Figure S1** - **Structural ensembles and target data used for HDX ensemble refinement testing**

**Figure S2** - **Residues included in target TeaA HDX data at reduced levels of sequence coverage**

**Figure S3** - **Root mean square deviation (RMSD) of TeaA $C_\alpha$ atoms across the unbiased trajectory from bias-exchange metadynamics simulations**

**Figure S4** - **Relationship between $\gamma$, $W_{app}$, and agreement with target data in HDX reweighting analyses**

**Figure S5** - **Structural variance in the clustered ensembles of TeaA after ensemble reweighting**

**Figure S6** - **Effect of overlap in the peptide segment residue ranges on reweighting**

**Movie S1** - **Artificially-generated morph between the closed and open representative TeaA structures**

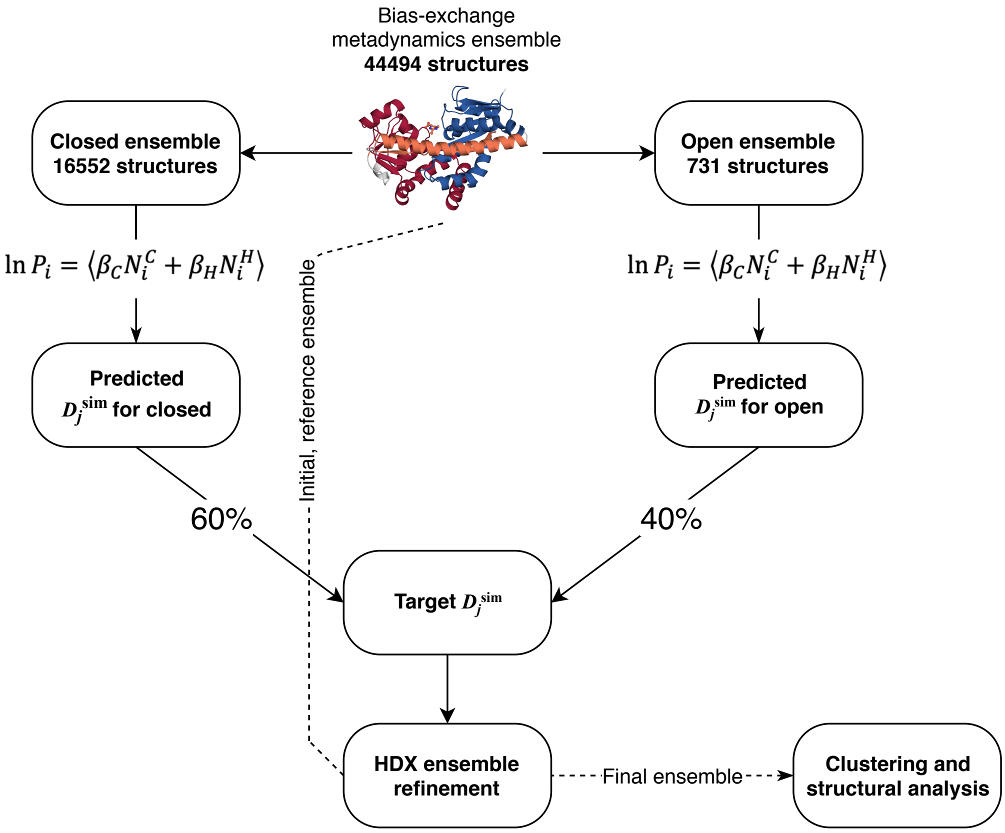


**Figure S1** – Structural ensembles and target data used for HDX ensemble refinement testing. Artificial HDX data (solid arrows) were generated for each exchangeable amide for a mixed ensemble corresponding to 60% closed and 40% open TeaA. These data were used for HDX ensemble refinement of the full test ensemble including closed, open, and decoy semi-open frames (dashed lines). Target data was optionally degraded so that deuterated fractions were averaged over peptide segments, or at a reduced level of sequence coverage (see main text Methods).

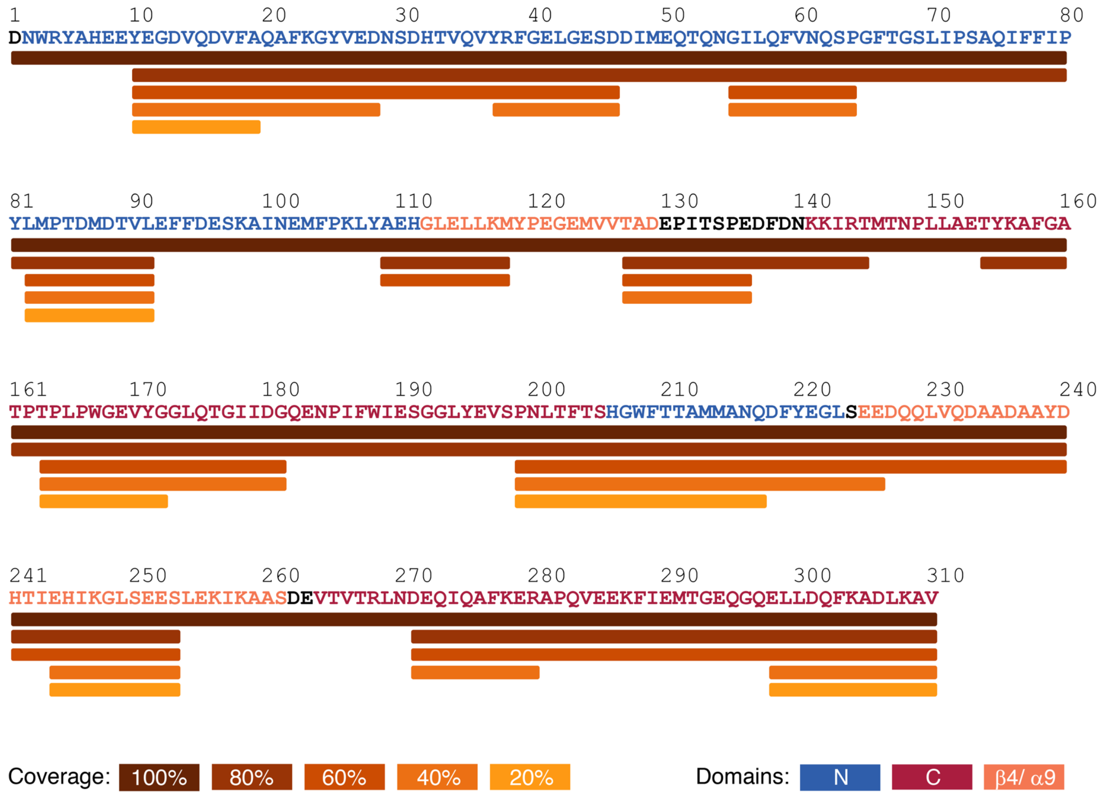


**Figure S2** – Residues included in target TeaA HDX data with reduced levels of sequence coverage. The sequence is colored according to the protein domain region as defined in Fig. 1A. Bars underneath the sequence indicate the residues included in the target data at 100% coverage (dark brown) and at approximately 80, 60, 40, or 20% coverage (from light brown to orange to yellow). See main text Methods for more details.

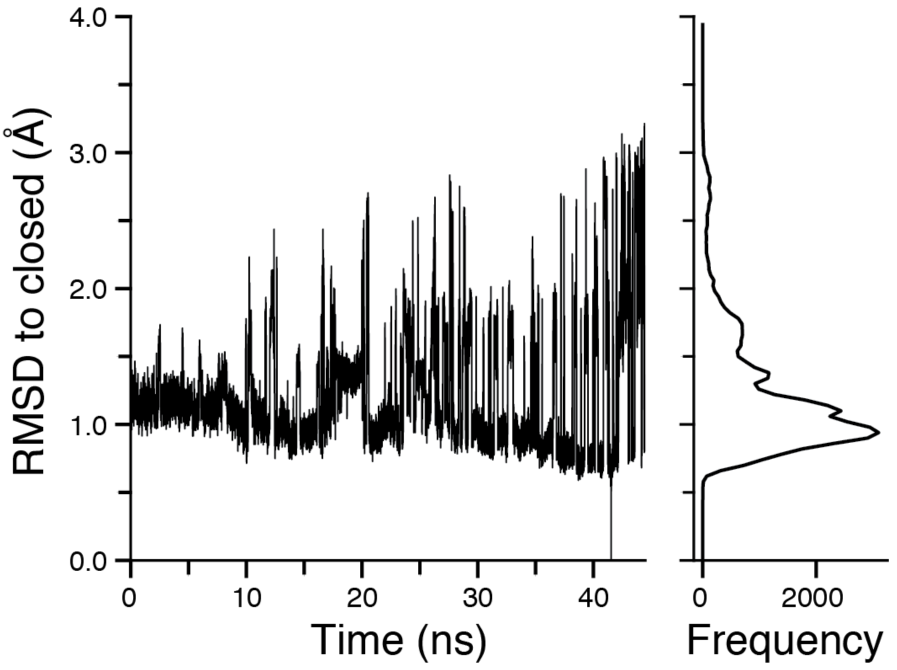


**Figure S3** – Structural variability in the reference simulation of TeaA and selection of reference structures. The root mean square deviation (RMSD) of the $C_\alpha$ atoms from the reference closed conformation is plotted against time for the unbiased trajectory from bias-exchange metadynamics simulations performed by Marinelli & coworkers (1). Trajectory frames were taken at 1 ps intervals. The frame at 41.546 ns was arbitrarily selected as the reference closed configuration. The frame at 44.428 ns has the highest RMSD to the closed conformation and was therefore chosen as the reference open configuration. The distribution of the RMSD values to the closed structure is shown in the panel on the right.

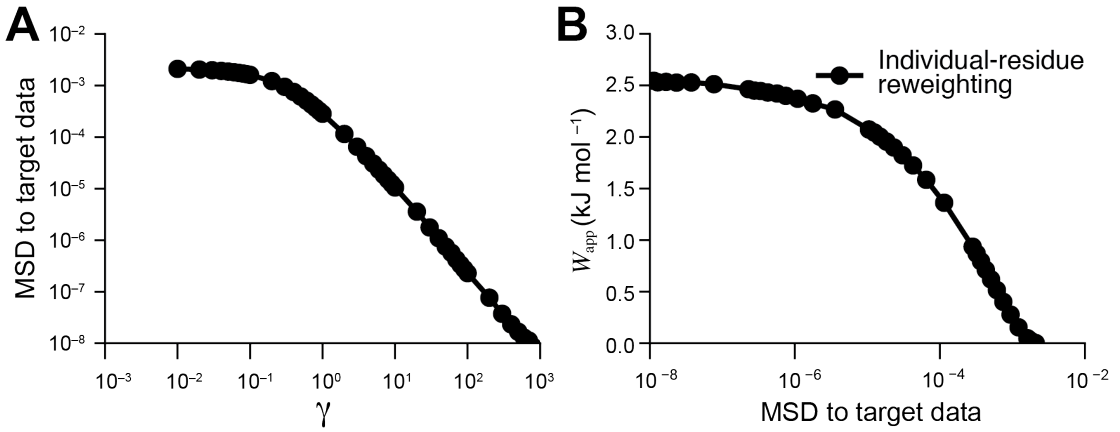


**Figure S4** – Relationship between γ, $W_{\text{app}}$, and the agreement with the target data during HDX reweighting analyses targeting TeaA HDX data with residue-level resolution. (**A**) The mean square deviation (MSD) between the predicted and target HDX values decreases as the value of γ increases. Little improvement in MSD is observed below $\gamma \approx 10^{-1}$, suggesting that, beyond this point, the initial HDX data lies within the uncertainty distribution $\rho_{\text{err}}$ defined by γ. (**B**) For the same reweighting analyses, the reduction in MSD is coupled to an increase in $W_{\text{app}}$ until a plateau is reached for MSD values below ca. $10^{-7}$.

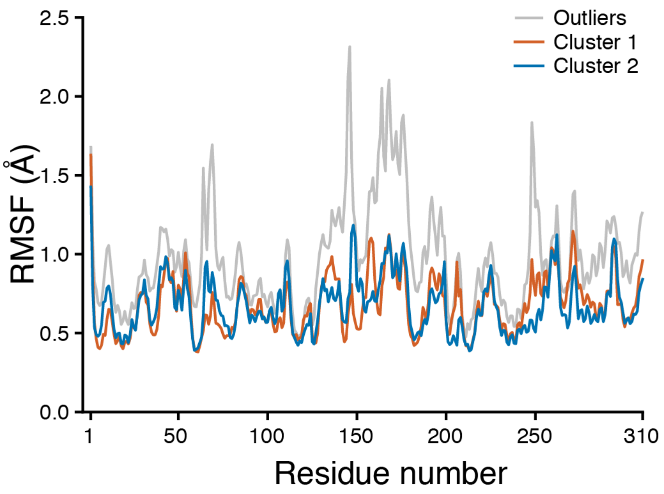


**Figure S5** – Structural variance in the clustered ensembles of TeaA after ensemble reweighting by residue-resolved HDX data. The backbone root mean squared fluctuation (RMSF) was averaged over each residue for all conformations in the two main clusters (orange and cyan), or in the outliers (gray).

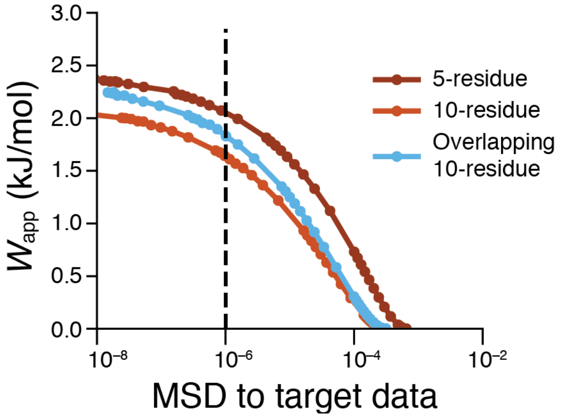


**Figure S6** – Effect of overlap (i.e. redundancy) in the peptide segment residue ranges on reweighting. Decision plot showing the work applied during reweighting, against the MSD of the reweighted ensemble to target HDX data. Circles indicate independent reweighting experiments. Target data with overlapping 10-residue segments were defined such that peptides maintained 100% sequence coverage, but overlapped in approximately 5-residue intervals (e.g. residues 1-10, 6-15, 10-19, 15-24…). Reweighting using overlapping segments (blue) initially shows MSD similar performance to reweighting with non-overlapping 10-residue segment data (light brown), but trends towards the results obtained with shorter, 5-residue, peptide segments (dark brown), owing to the additional information content of target data with peptide redundancy.

**Movie S1** – Artificially-generated morph between the closed and open representative structures of TeaA. TeaA is shown in cartoon representation (wheat), and the ectoine substrate from the closed configuration shown in ball and stick representation (peach).
